# Smaller Hippocampal CA-1 Subfield Volume in Posttraumatic Stress Disorder

**DOI:** 10.1101/337030

**Authors:** Lyon W. Chen, Delin Sun, Sarah L. Davis, Courtney C. Haswell, Emily L. Dennis, Chelsea A. Swanson, Christopher D. Whelan, Boris Gutman, Neda Jahanshad, Juan Eugenio Iglesias, Paul Thompson, Mid-Atlantic MIRECC Workgroup, H. Ryan Wagner, Philipp Saemann, Kevin S. LaBar, Rajendra A. Morey

## Abstract

**Background:** Smaller hippocampal volume in patients with PTSD represents the most consistently reported structural alteration in the brain. Subfields of the hippocampus play distinct roles in encoding and processing of memories, which are disrupted in PTSD. We examined PTSD-associated alterations in 12 hippocampal subfields in relation to global hippocampal shape, and clinical features.

**Methods:** Case-control cross-sectional study of US military veterans (n=282) from the Iraq and Afghanistan era were grouped into PTSD (n=142) and trauma-exposed controls (n=140). Participants underwent clinical evaluation for PTSD and associated clinical parameters followed by MRI at 3-Tesla. Segmentation with FreeSurfer v6.0 produced hippocampal subfield volumes for the left and right CA1, CA3, CA4, DG, fimbria, fissure, hippocampus-amygdala transition area, molecular layer, parasubiculum, presubiculum, subiculum, and tail, as well as hippocampal meshes. Covariates included age, gender, trauma exposure, alcohol use, depressive symptoms, antidepressant medication use, total hippocampal volume, and MRI scanner model.

**Results:** Significantly lower subfield volumes were associated with PTSD in left CA1 (*p*=.01; *d*=.21; uncorrected), CA3 (*p*=.04; *d*=.08; uncorrected), and right CA3 (*p*=.02; *d*=.07; uncorrected) only if ipsilateral whole hippocampal volume was included as a covariate. A trend level association of L-CA1 with PTSD [F_4, 221_=3.32, p = 0.07] is present and the other subfield findings are non-significant if ipsilateral whole hippocampal volume is not included as a covariate. PTSD associated differences in global hippocampal shape were non-significant.

**Conclusions:** The present finding of smaller hippocampal CA1 in PTSD is consistent with model systems in rodents that exhibit increased anxiety-like behavior from repeated exposure to acute stress. Behavioral correlations with hippocampal subfield volume differences in PTSD will elucidate their relevance to PTSD, particularly behaviors of associative fear learning, extinction training, and formation of false memories.

## INTRODUCTION

Individuals with posttraumatic stress disorder (PTSD) may experience deficits in declarative memory such as remembering events, facts or lists, fragmentation of autobiographical or trauma-related memories, and trauma-related amnesia [1]. The hippocampus plays an important role in memory formation and retrieval that has long been implicated in the clinical presentation of PTSD [2]. Indeed, lower hippocampal volume in PTSD has been a reliably reported structural alteration for over two decades [3; 4]. We sought to attain improved spatial and functional characterization of this structural alteration via two complimentary approaches, (1) quantify the volume of 12 hippocampal subfields, and (2) conduct 3-D vertex-based shape analysis of the hippocampus to identify localized surface features associated with PTSD.

Careful investigation of PTSD-associated alterations in hippocampal subfields and their relationship to differences in hippocampal shape, and clinical features is scant, has met with inconsistent results, and has been beset by limitations [5-7]. Employing manual subfield segmentation, Wang et al reported lower volume in CA3 associated with PTSD, Hayes et al found smaller dentate gyrus (DG), whereas Mueller et al found PTSD associated no subfield differences. The primary limitation of previous studies was small sample size of n=36 given the expected range of effect sizes. Hayes et al studied the largest sample to date (n=97), but used subfield segmentation with FreeSurfer v5.3 that suffered from three major shortcomings [8]. First, the resolution of the *in vivo* training data was insufficient for the human raters to accurately distinguish subregions, forcing excessive reliance on geometric boundary criteria for tracing subfields, but on the other hand may be able to overcome image artifacts sometimes overlooked by automated techniques. The second issue was that the delineation protocol was designed for the hippocampal body, which translated poorly to the hippocampal head and tail. The resultant third problem was that the volumes of the subregions did not agree well with histological studies. Therefore, FreeSurfer v5.3 is based on an anatomically incorrect atlas [9], notably for CA1. These shortcomings are addressed in the present study by a completely new atlas in FreeSurfer v6.0 that was built with a novel atlasing algorithm and *ex vivo* MRI data acquired from *post mortem* brains [10]. Thus, FreeSurfer v6.0 combines *ex-vivo* and *in-vivo* scans, with the former acquired on 15 *ex-vivo* brains scanned at 7-Tesla to attain extremely high signal to noise ration and 130-µm^3^ isotropic resolution, whereas v5.3 used *in vivo* atlas from five cases acquired at nearly 9-fold lower resolution of 380-µm^3^ resolution and the manual segmentation studies were acquired at 4T with 180-fold lower resolution of 400 ×; 500 µm in-plane and 2000-µm through-plane resolution. These enhancements enable segmentation of 12 subfields with FreeSurfer v6.0 as compared to five subfields with v5.3 or manual segmentation methods.

Subfields of the hippocampus are involved in discrete aspects of memory encoding and consolidation. For instance, the dentate gyrus (DG) is important in distinguishing features that are different from other memories in order to store similar memories as discrete events – a phenomenon called *pattern separation* [11; 12]. Pattern separation deficits may underlie fear generalization [13], a process that occurs in anxiety and stress based disorders including PTSD [14]. By contrast, the entorhinal cortex (EC) and *cornu ammonis* subfield-3 (CA3) are crucial in recognizing different events with overlapping features – a phenomenon called *pattern completion* that has important implications in contextual fear conditioning [15] and is a widely investigated model of PTSD [16]. Chronic stress in rats produces atrophy and debranching of dendrites in pyramidal neurons of the CA3 [17] and decreased neurogenesis in the DG [18], which appear to be reversible in these models when stress is alleviated. Preclinical research in rats has shown that the CA1 subfield is involved in context-specific memory retrieval after extinction [19]. These and other findings clearly point to a major influence of hippocampal CA1 neurons in conditioned fear and its extinction [20]. Extinction learning, which relies critically on intact CA1 function [20; 23; 24], is impaired in widely adopted experimental models of PTSD and consistent with re-experiencing symptoms of PTSD.

The capability for automated, *in vivo* segmentation of the human hippocampal subfields is now imperative for achieving replicability and reproducibility efficiently in large multi-site initiatives [25; 26]. The recently released FreeSurfer v6.0 makes it possible to estimate the hippocampal subfields from 1-mm T1-weighted MRI. Despite the fact that historically, segmentation of subfields was usually based on higher resolution images, that may include T2, with 0.2-0.7 mm and limited contrast between some of the subfields at 1-mm resolution, it has been shown that segmentations estimated from 1-mm scan resolution nevertheless carry useful information on subfield volumes [10]. The segmentation algorithm was validated on three publicly available datasets with varying MRI contrast and resolution [27].

Our goal was to investigate the association between PTSD and 12 hippocampal subfield volumes, as well as the relationship of subfield volume to hippocampal shape among a large cohort of younger US military veterans. Based on the foregoing evidence from animal and human research, we hypothesized the PTSD group would have smaller CA1, CA3, and DG volumes. Given the application of an anatomically incorrect atlas for CA1 in FreeSurfer v5.3 that was corrected in FreeSurfer v6.0, and the unique role of CA1 in context-specific memory retrieval, a strongly implicated behavioral deficit in PTSD, we elevated our prediction probability of smaller CA1 volume relative to CA3 and DG. Furthermore, a complimentary analysis of hippocampal shape was expected to reveal differences for surfaces corresponding to the affected subfields.

## METHODS

### Participants

We enrolled a total of 290 Iraq and Afghanistan era military service veterans, which were recruited from our repository [28]. Among these participants, 282 were selected for analysis following quality control (QC) procedures that consisted of 142 individuals with PTSD and 140 trauma exposed controls. Eight scans failed segmentation QC (non-hippocampal tissue assigned to CA1, contrast insufficient to identify the hippocampal fissure, presence of holes) resulting in participant exclusion. Participants were screened for inclusion/exclusion criteria based on information available in the repository. Important exclusions included major Axis I diagnosis (other than MDD or PTSD), contraindication to MRI, moderate/severe traumatic brain injury, substance dependence, neurological disorders, and age over 65 years. All participants provided written informed consent to participate in procedures reviewed and approved by the Institutional Review Boards at Duke University and the Durham VA Medical Center. Participants’ age, sex, and other demographic and clinical information are summarized in **Table 1** (see Supplement).

**Table 1:**
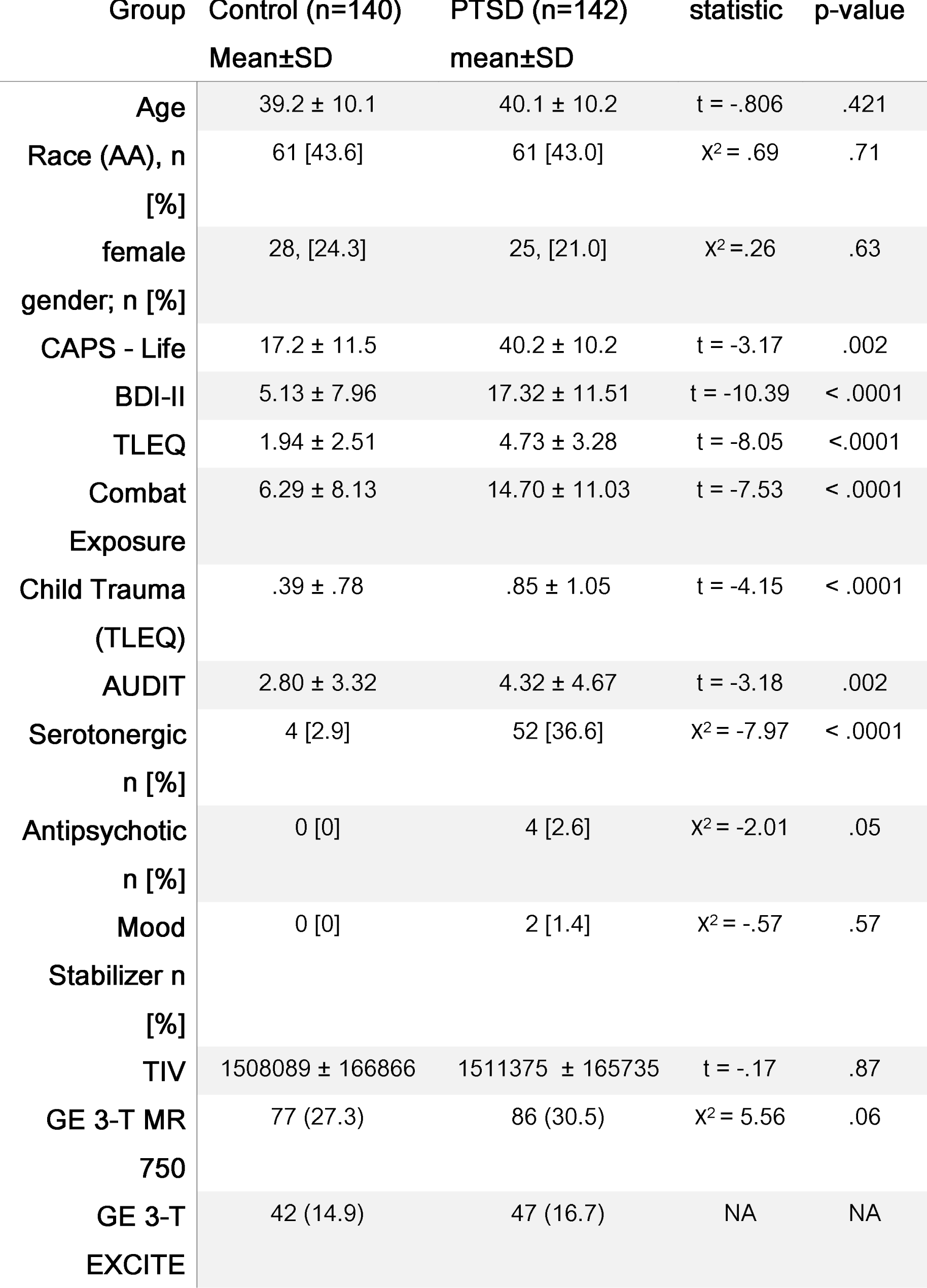
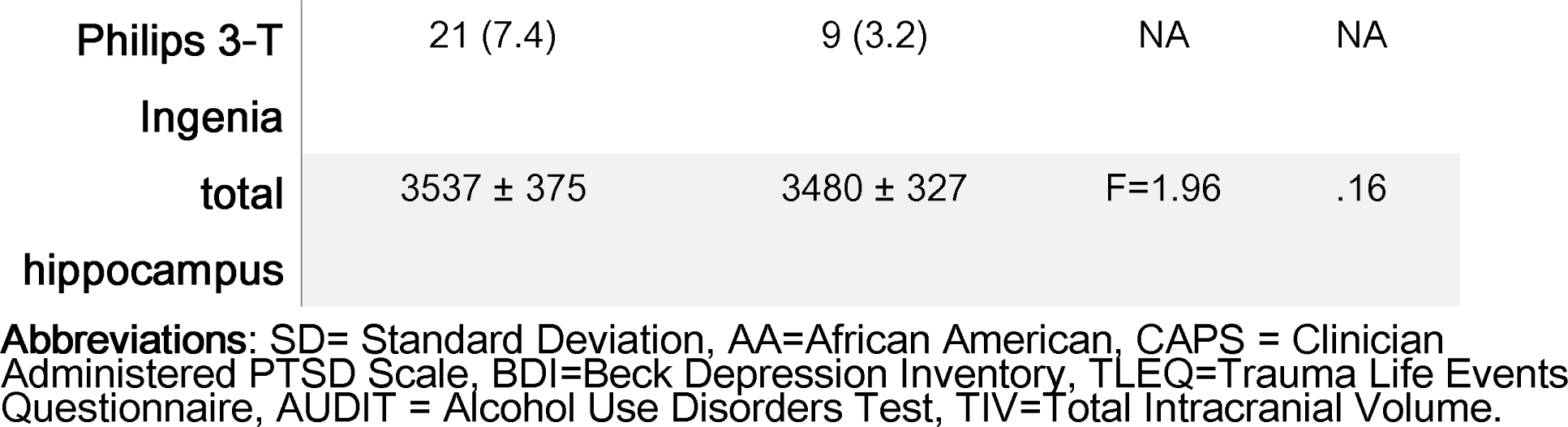
Clinical and Demographic Features of Sample.

| Group | Control (n=140)<br>Mean±SD | PTSD (n=142)<br>mean±SD | statistic | p-value |
| --- | --- | --- | --- | --- |
| Age | 39.2 ± 10.1 | 40.1 ± 10.2 | t = -.806 | .421 |
| Race (AA), n [%] | 61 [43.6] | 61 [43.0] | $\chi^2 = .69$ | .71 |
| female gender; n [%] | 28, [24.3] | 25, [21.0] | $\chi^2 = .26$ | .63 |
| CAPS - Life | 17.2 ± 11.5 | 40.2 ± 10.2 | t = -3.17 | .002 |
| BDI-II | 5.13 ± 7.96 | 17.32 ± 11.51 | t = -10.39 | < .0001 |
| TLEQ | 1.94 ± 2.51 | 4.73 ± 3.28 | t = -8.05 | <.0001 |
| Combat Exposure | 6.29 ± 8.13 | 14.70 ± 11.03 | t = -7.53 | < .0001 |
| Child Trauma (TLEQ) | .39 ± .78 | .85 ± 1.05 | t = -4.15 | < .0001 |
| AUDIT | 2.80 ± 3.32 | 4.32 ± 4.67 | t = -3.18 | .002 |
| Serotonergic n [%] | 4 [2.9] | 52 [36.6] | $\chi^2 = -7.97$ | < .0001 |
| Antipsychotic n [%] | 0 [0] | 4 [2.6] | $\chi^2 = -2.01$ | .05 |
| Mood Stabilizer n [%] | 0 [0] | 2 [1.4] | $\chi^2 = -.57$ | .57 |
| TIV | 1508089 ± 166866 | 1511375 ± 165735 | t = -.17 | .87 |
| GE 3-T MR 750 | 77 (27.3) | 86 (30.5) | $\chi^2 = 5.56$ | .06 |
| GE 3-T EXCITE | 42 (14.9) | 47 (16.7) | NA | NA |
| **Philips 3-T** | 21 (7.4) | 9 (3.2) | NA | NA |
| **Ingenia** |  |  |  |  |
| **total** | 3537 ± 375 | 3480 ± 327 | F=1.96 | .16 |
| **hippocampus** |  |  |  |  |

**Abbreviations:** SD= Standard Deviation, AA=African American, CAPS = Clinician Administered PTSD Scale, BDI=Beck Depression Inventory, TLEQ=Trauma Life Events Questionnaire, AUDIT = Alcohol Use Disorders Test, TIV=Total Intracranial Volume.

### MRI acquisition

All images were acquired on a 3-Tesla scanner equipped with an 8-channel head coil. All scans were acquired as high-resolution T1-weighted whole-brain axial images with 1-mm isotropic voxels on three different scanners: [1] GE Discovery MR750 (n=163, 57.8%; Control n=77, PTSD n=86) [2] GE Signa EXCITE (n=89, 31.5%; Control n=42, PTSD n=47), and [3] Philips Ingenia scanner (n=30, 10.6%; Control n=21, PTSD n=9). See Online Supplement for ‘Image Acquisition Parameters’.

### Hippocampal Subfield Volume Analysis

Automated segmentation and labeling of subcortical volumes and estimation of total intracranial volume (TIV) from T1 images was performed using the FreeSurfer image analysis suite[29] (v5.3.0; http://surfer.nmr.mgh.harvard.edu/) and its library tool, *recon-all*. Hippocampal subfield segmentation was performed using FreeSurfer 6.0 and its library function, *hippocampal-subfields-T1*. Hippocampal subfield volumes for the left and right hemispheres were generated in each subject for the CA1, CA3, CA4, DG, fimbria, fissure, hippocampus-amygdala transition area (HATA), molecular layer, parasubiculum, presubiculum, subiculum, and tail (**Figure 1**). We applied standardized protocols for image analysis for subcortical segmentation developed by the Consortium for <u>EN</u>hancing Imaging <u>G</u>enetics through <u>M</u>eta-<u>A</u>nalysis (ENIGMA; see Supplement), (http://enigma.ini.usc.edu/protocols/imaging-protocols/). We have previously analyzed FreeSurfer output from multiple scanners with evidence that results are robust to this heterogeneity. In Logue et al we used 16 different scanners and found *p*_het_=0.74 and I^2^=0 for hippocampal volume. In Whelan et al [25], we demonstrated high intraclass correlation (ICC) across 1.5T and 3.0-T scanners with FreeSurfer v6.0 for hippocampal subfields including CA1 (ICC = 0.915) and CA3 (ICC=0.827).

**Figure 1:**
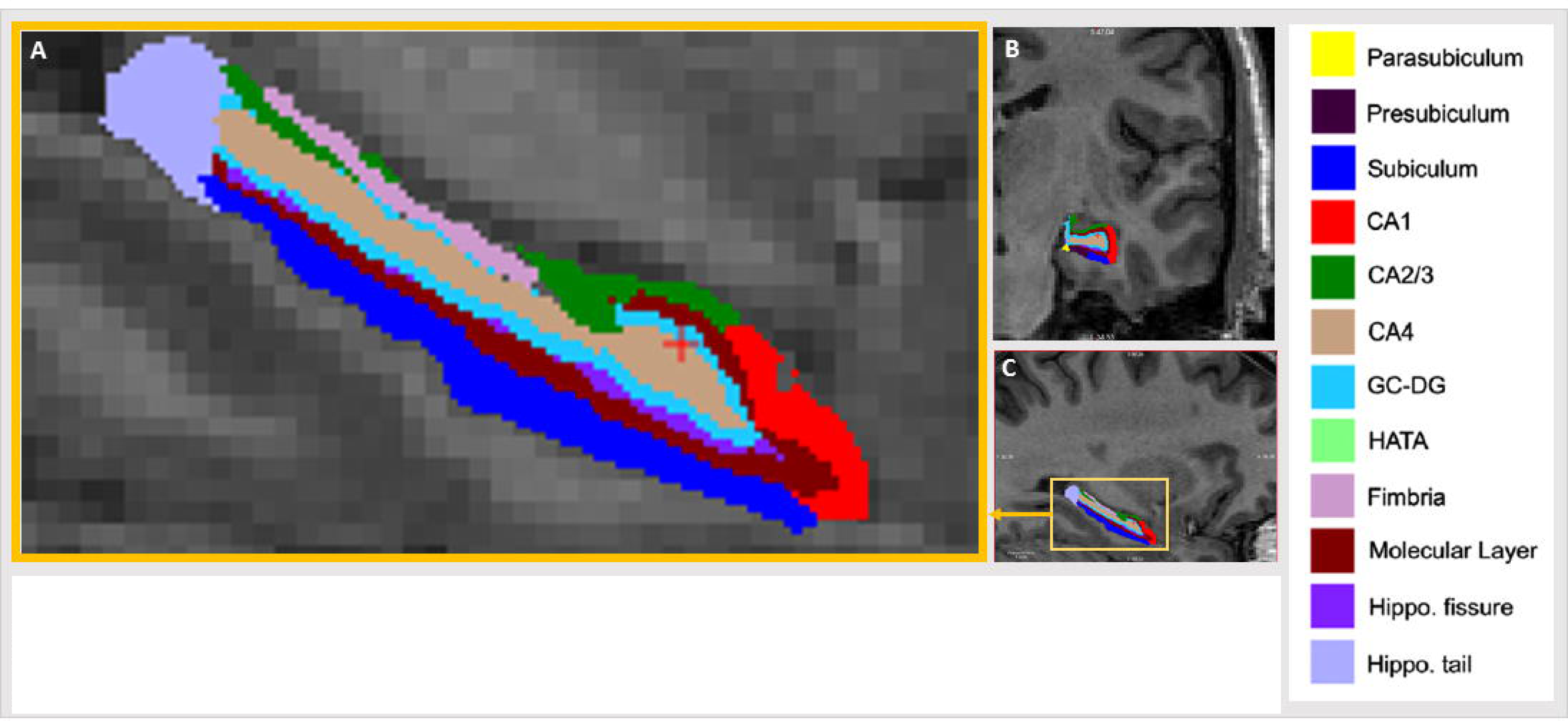
Hippocampal Subfield Segmentation. Automated segmentation of the hippocampus into 12 subfields in each hemisphere of the brain was performed with FreeSurfer v6.0. Subfield images of CA1, CA2/3, CA4, dentate gyrus, hippocampal-amygdala transition area (HATA), subiculum, tail, fissure, presubiculum, parasubiculum, molecular layer, fimbria are shown in **(a)** magnified sagittal, **(b)** coronal, and **(c)** sagittal planes.

### Quality Control Procedures

We applied quality assurance for hippocampal subfield segmentations using protocols developed by the ENIGMA-MDD Consortium and the ENIGMA-MDD hippocampal subfields project (see https://pgc-ptsd.com/wp-content/uploads/2017/08/PTSD_Instructions_Subfields_part_IR_II.pdf).

### Hippocampal Shape Analysis

We applied a standard analysis pipeline for subcortical shape developed by ENIGMA [30]. FreeSurfer segmentation and labels created from the volumetric analysis described above were used to generate meshes and shape data for the hippocampus. Vertex information from each subject was extracted to carry out between group analyses with regressors. We applied vertex-wide FDR correction based on 2,502 vertices (see Supplement).

### Statistical Analysis

The subfield volumes obtained from FreeSurfer was the dependent variable in an ordinary least square (OLS) regression model run separately for each subfield. Bonferroni correction for multiple testing was applied to the volumetric analyses given that 12 regions from 2 hemispheres were assessed. We included a covariate for ipsilateral whole hippocampal volume because CA1 volume is highly correlated with whole hippocampal volume, i.e. individuals with smaller hippocampi will have smaller CA1 in much the same way that individuals with a small brain (TIV) will have a small hippocampus. Further details of the regression model and regressors are in the **Supplement**. Results for the CA1, CA3, and DG were not Bonferroni corrected because of prior evidence of volume reduction reported in adults exposed to childhood maltreatment [31] and adult PTSD [6; 7]. Regressors in the initial analysis were selected based on established associations with hippocampal volume, but subsequent re-analysis included only regressors with *p* < 0.15 (see **Supplement**).

The preexisting group difference in the level of trauma exposure between the PTSD and Control groups meant that group differences could be attributable to either PTSD or trauma exposure. Inclusion of trauma exposure as covariate could lead to inconclusive findings because removing the variance associated with trauma exposure will also alter variance in the dependent variable associated with diagnostic groups as explained by Miller and Chapman [32]. The role of trauma exposure on hippocampal volume was further scrutinized vis-à-vis PTSD with correlations between trauma exposure and each of the significant subfields results in PTSD and Control groups. The difference in correlations between the PTSD and Control group was compared with Fisher’s r-to-z test [33].

## RESULTS

### Demographic and clinical characteristics

In broad terms, exposure to combat trauma and lifetime trauma, as well as symptoms of depression and alcohol use were significantly greater in the PTSD group than the trauma-exposed control group. Detailed clinical and demographic information is reported by diagnostic group in **Table 1**.

### Association of subfield volumes with PTSD

We found PTSD was associated with significantly lower volume in L-CA1 (*p*=.01; Cohen’s *d*=.21), L-CA3 (*p*=.04; Cohen’s *d*=.08), and R-CA3 (*p*=.02; Cohen’s *d*=.07). The data was reanalyzed for regressors with *p* < 0.15 in the initial univariate model. These results that were fully consistent with the initial analyses with all regressors as listed in **Table 2** and the **Supplement**. The largest effect size among regions with significant between-group differences was in L-CA1 (Cohen’s *d* = .21), whereas the effect size was small for L-CA3 and R-CA3.

**Table 2:**
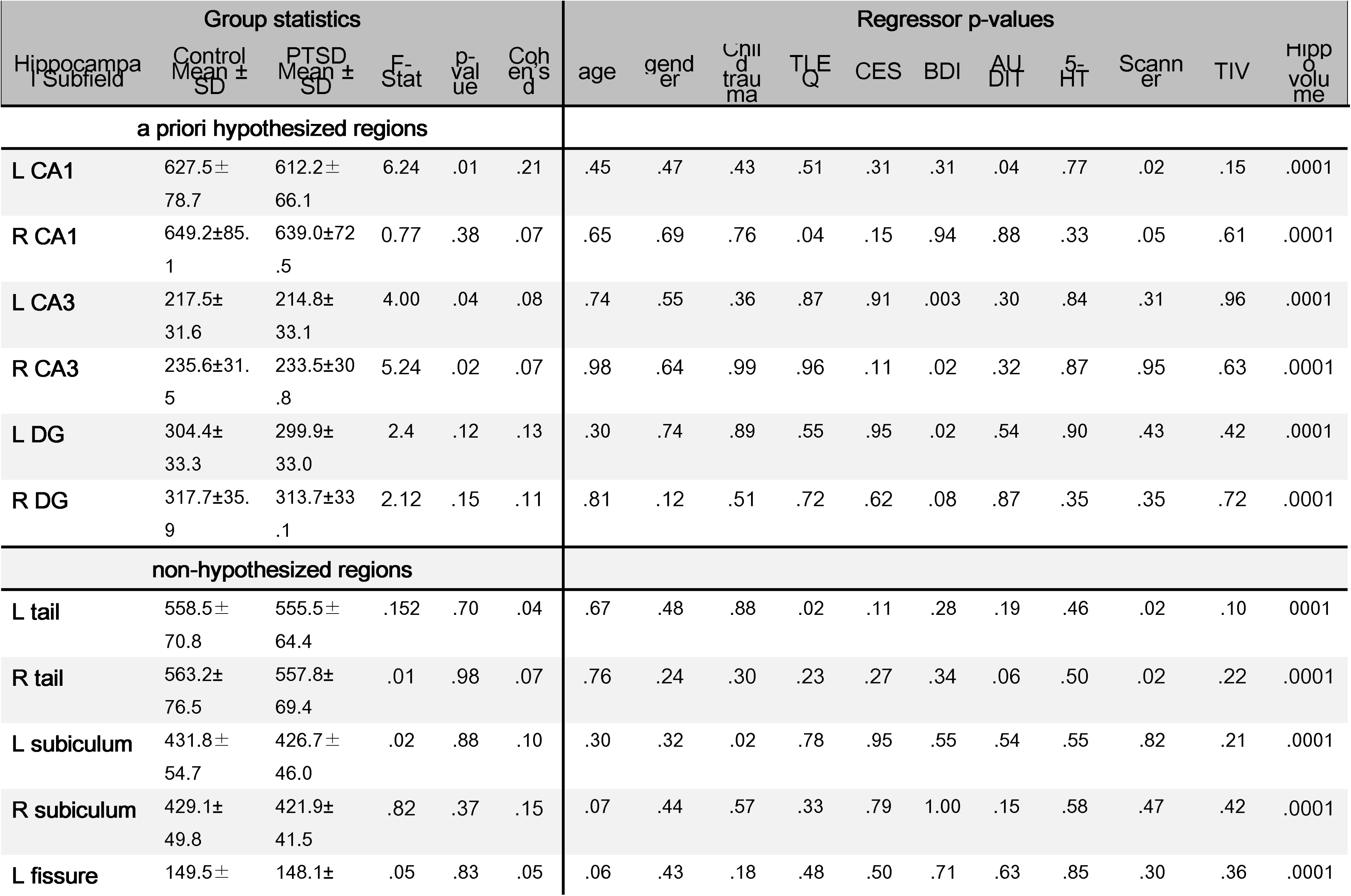
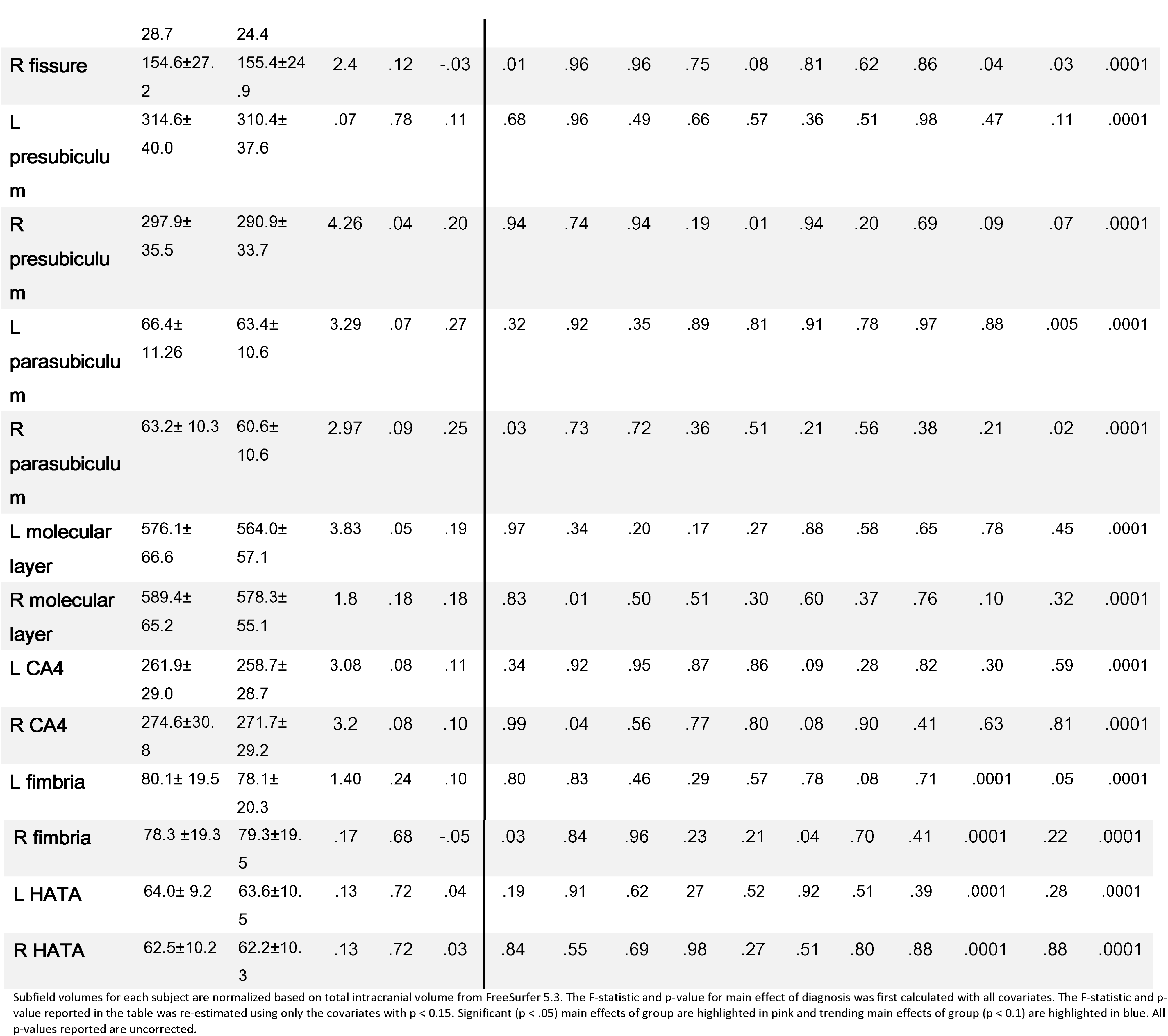
Hippocampal Subfield Volume Association with PTSD.

There were no other subfields with significant between-group differences after correction for multiple comparisons. Detailed results of descriptive and inferential statistics including the role of each regressor are provided in **Table 2**. Serotonergic medication usage, which was a regressor in the initial analyses, was non-significant for all subfields. Results of secondary analysis that omitted data of subjects on antipsychotic medication (n=4) yielded results that were consistent with the main analyses, L-CA1 [F_4, 273_=5.76, p=0.017], L-CA3 [F_4, 273_=4.27, p=0.04], and at trend level for R-CA3 [F_4, 273_=3.80, p=.052]. To address the possible role of SSRI medication, we repeated main analyses by excluding subjects taking SSRI medication (n=56). Our results were consistent with our main findings in the L-CA1 [F_4, 220_=4.76, p=0.03], L-CA3 [F_4,_ 220 = 3.80, p=0.052], and at trend level for R-CA3 [F_4, 220_=3.33, p=.07]. Among regressors, the whole hippocampal volume regressor (ipsilateral) showed the most consistent results. Subfields with very small volumes (≤ 100 mm^3^) consistently demonstrated significant effect of scanner type. In addition, the gender regressor for R-CA4, combat exposure for R-presubiculum, depression symptoms regressor for L-DG, L-CA3, R-CA3, and R-fimbria were nominally significant (**Table 2)**.

Excluding ipsilateral whole hippocampal volume as a covariate resulted in a trend level association of L-CA1 with PTSD [F_4, 221_=3.32, p = 0.07] and non-significant results for the L-CA3 [F_4, 221_=0.07, p = 0.79] and R-CA3 [F_4, 221_=0.23, p = 0.63] (**Table 3**). To understand the role of including ipsilateral whole hippocampal volume as a covariate, we plotted the L-CA1, L-CA3, and R-CA3 residualized values with the covariates included in the regression model, importantly the ipsilateral whole hippocampal volume (**Figure 2**).

**Table 3:**
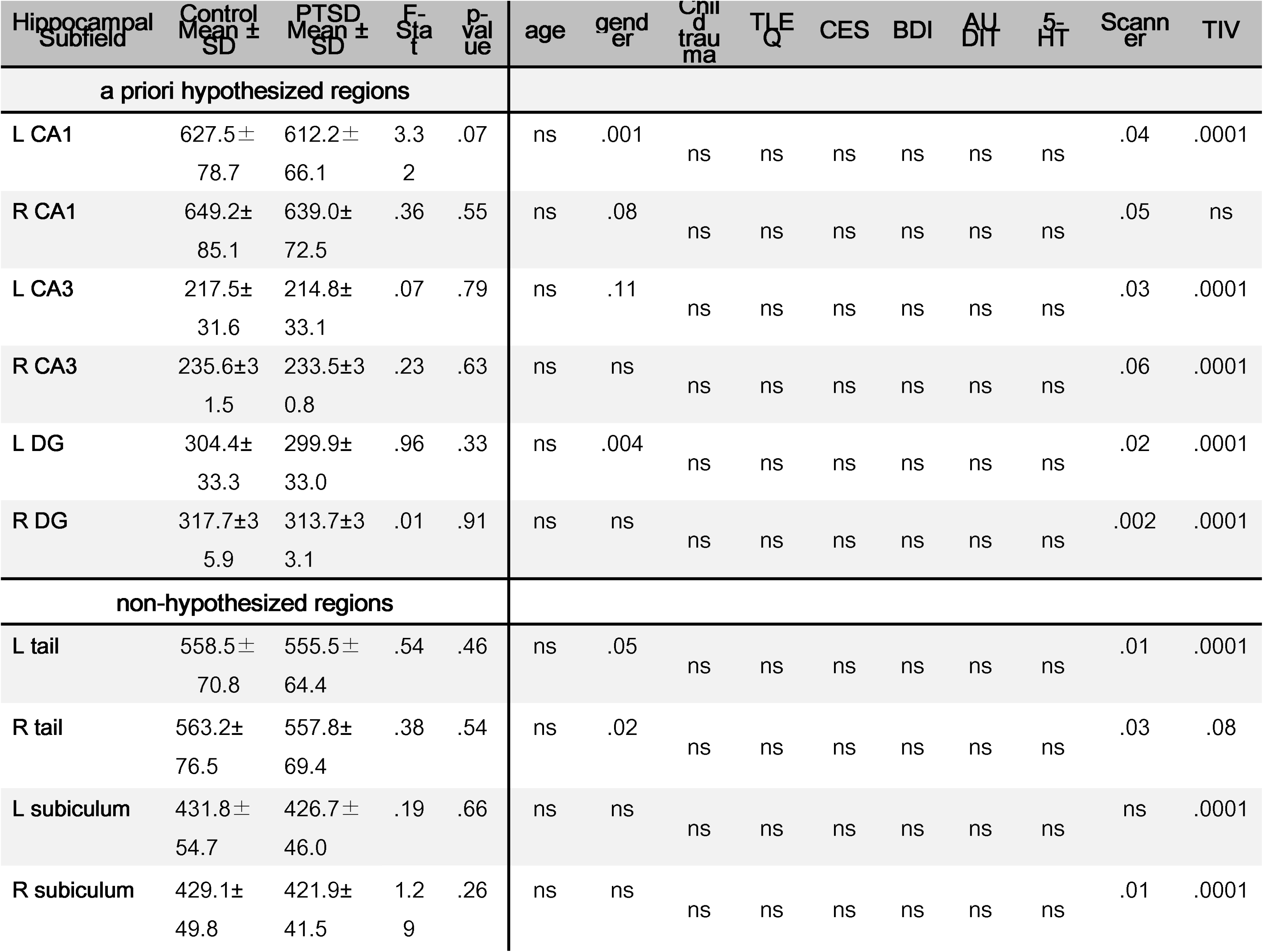
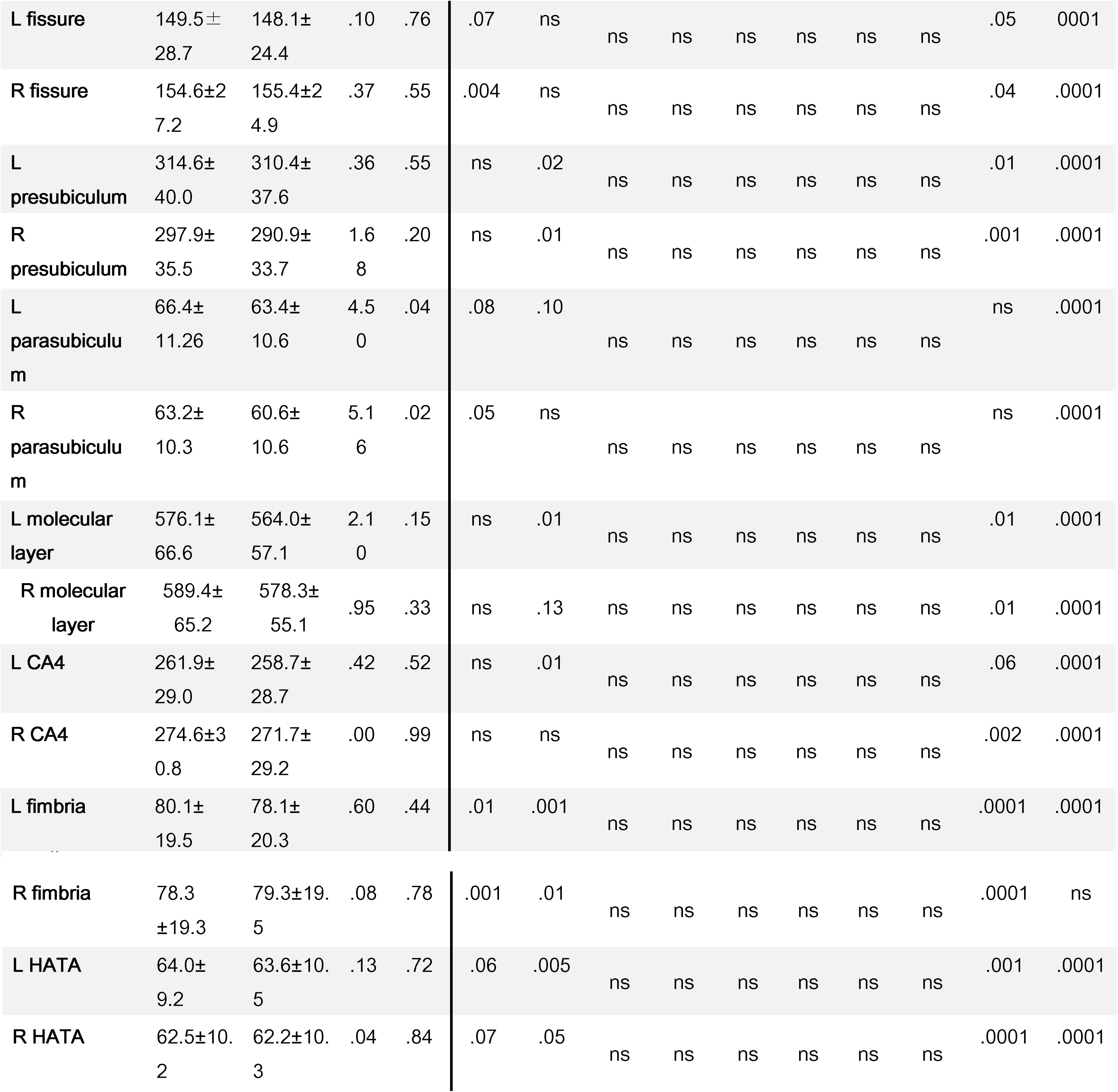
Hippocampal Subfield Volume Association with PTSD Excluding Ipsilateral Hippocampal Volume Covariate.

**Table 3:** Effects of Aging on Hypothesized Subfields.

| Group | Correlation (r) | p-value | Trauma Control (r) | Trauma Control p-value | PTSD (r) | PTSD p-value | Interaction F-statistic | Interaction p-value |
| --- | --- | --- | --- | --- | --- | --- | --- | --- |
| L-CA1 | -.10 | .08 | -.02 | .83 | -.19 | .03 | 1.78 | .18 |
| R-CA1 | -.11 | .07 | -.05 | .52 | -.16 | .05 | .56 | .46 |
| L-CA3 | -.08 | .18 | -.02 | .77 | -.13 | .13 | .87 | .35 |
| R-CA3 | -.05 | .41 | -.01 | .87 | -.08 | .32 | .33 | .57 |
| L-DG | -.13 | .04 | -.07 | .41 | -.17 | .04 | .85 | .36 |
| R-DG | -.08 | .18 | -.03 | .76 | -.13 | .11 | .73 | .39 |

Subfield volumes for each subject are normalized based on total intracranial volume from FreeSurfer 5.3.
Interaction tests for significantly different slopes of aging on subfield volume in PTSD vs. trauma-exposed control groups

**Figure 2:**
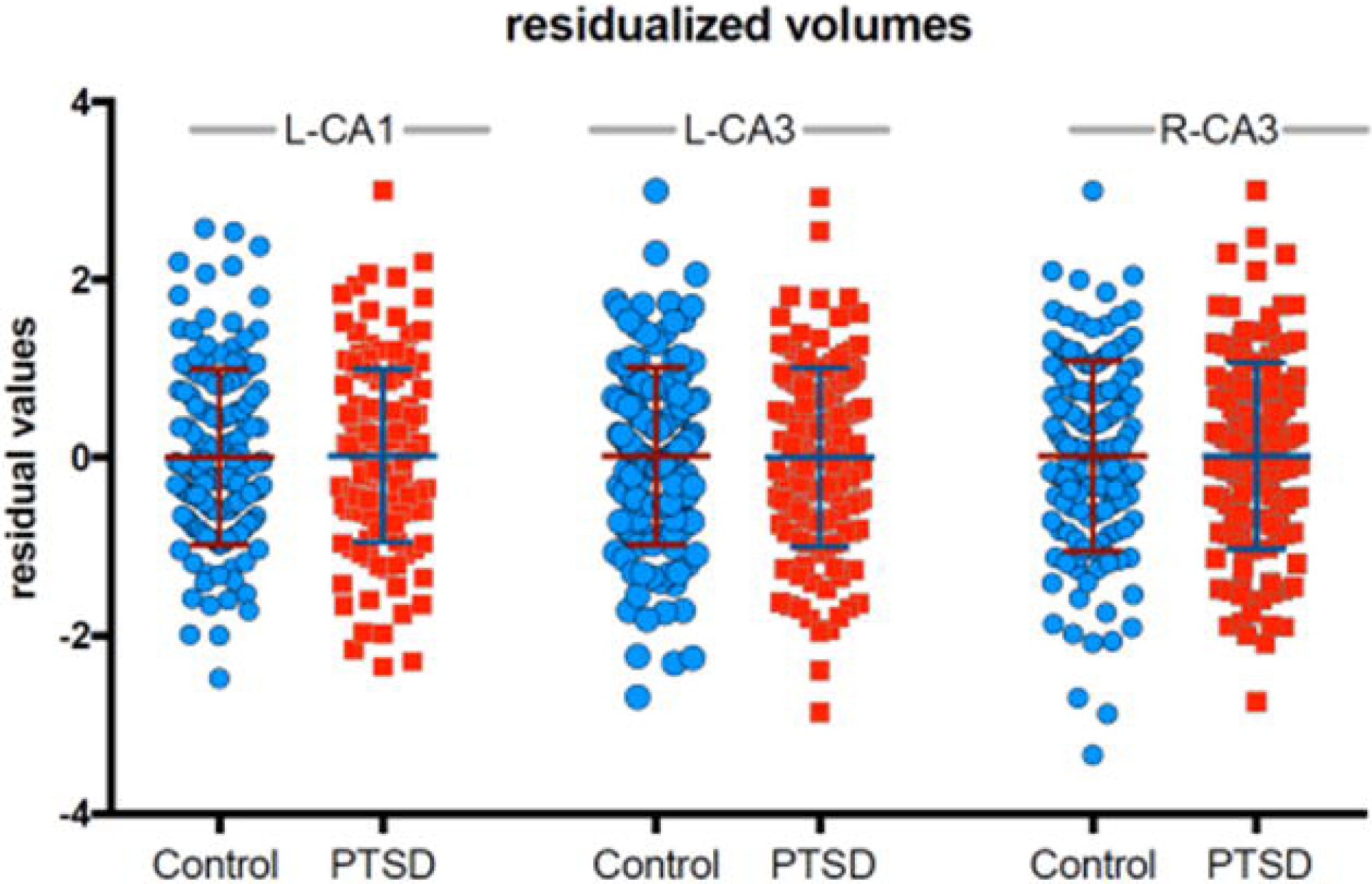
The L-CA1, L-CA3, and R-CA3 values that are residualized for the covariates included in the regression model, which includes the ipsilateral whole hippocampal volume covariate.

### Association of subfield volume with trauma exposure

The association between trauma exposure and subfield volumes for L-CA1, L-CA3, and R-CA3 was examined because trauma exposure was collinear with diagnostic groups, i.e. significantly greater in the PTSD than Control group [*t*_281_=-8.05; *p* < .0001). This raised the possibility that subfield volumes were related to trauma exposure rather than PTSD. Correlations between trauma exposure and subfield volumes were weak (r’s < .15). The correlation strength between trauma exposure and subfield volumes was not significantly different in PTSD than in Control groups based on Fisher’s r-to-z for L-CA1 (*z*=-.055; *p*=0.95), L-CA3 (*z*=1.27; *p*=0.20), and R-CA3 (*z*=1.15; *p*=0.25) (**Figure 3**).

**Figure 3:**
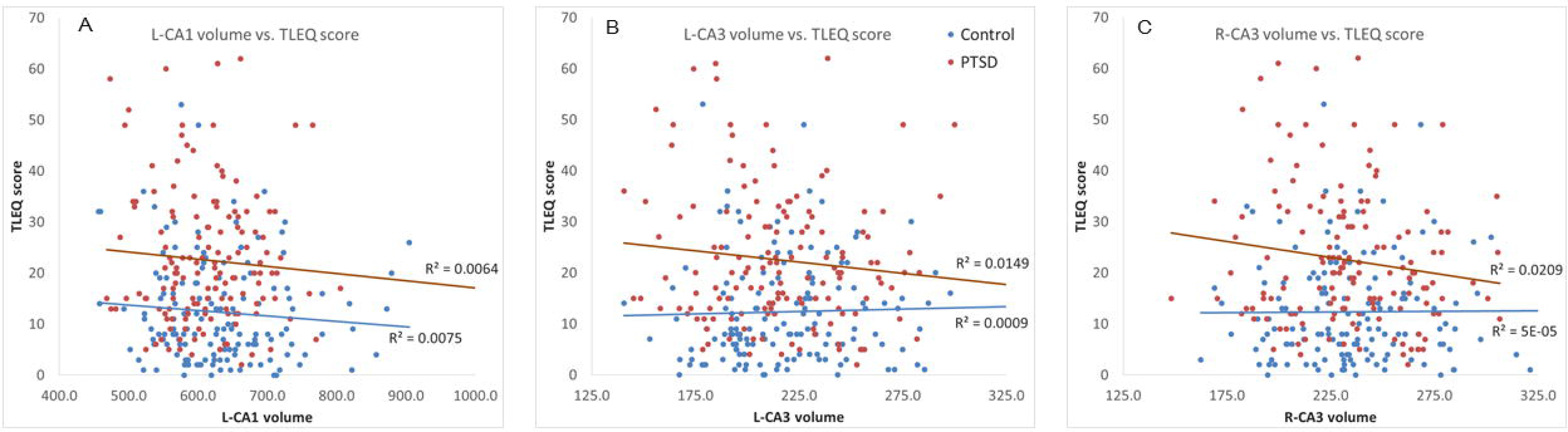
Results showed that the associations between lifetime trauma exposure (TLEQ score) and subfield volumes for **(A)** L-CA1, **(B)** L-CA3, and **(C)** R-CA3, were weak and were not significantly different in the PTSD group (red trend line) compared to the Control group (blue trend line).

The analysis that showed a significant association between R-CA3 and PTSD included a regressor for combat exposure based on a significant result for CES that was identified in the initial analysis, which included all regressors (**Table 2**). The CES score was significantly higher in the PTSD group than the control group [t_281_=-7.53; p<0.0001), which meant that the CES regressor was correlated with diagnostic grouping, again raising the possibility that the group-difference in R-CA3 was associated with combat exposure rather than PTSD. The correlation strengths between combat exposure and R-CA3 subfield volumes were weak (r’s < .06) and not significantly different in PTSD and Control groups using Fisher’s r-to-z (*z*=.79; *p*=0.43) [33].

### Effect of Age in PTSD

We found trend-level correlations between age and volume for bilateral CA1, CA2, and DG, but the interaction of age by PTSD diagnosis was non-significant (correlations did not differ significantly in the PTSD group relative to the trauma-exposed control group; **Table 3**) [6].

### Shape Results

The vertex-based analyses revealed that shape differences between the PTSD group and the control group were non-significant after vertex-wide FDR correction for multiple testing (p < .05). Results for trend-level significance with FDR correction (*p* < 0.2) are provided for β-map and *p*-map visualization of shape differences based on the Jacobian determinant in **Figure 4**.

**Figure 4:**
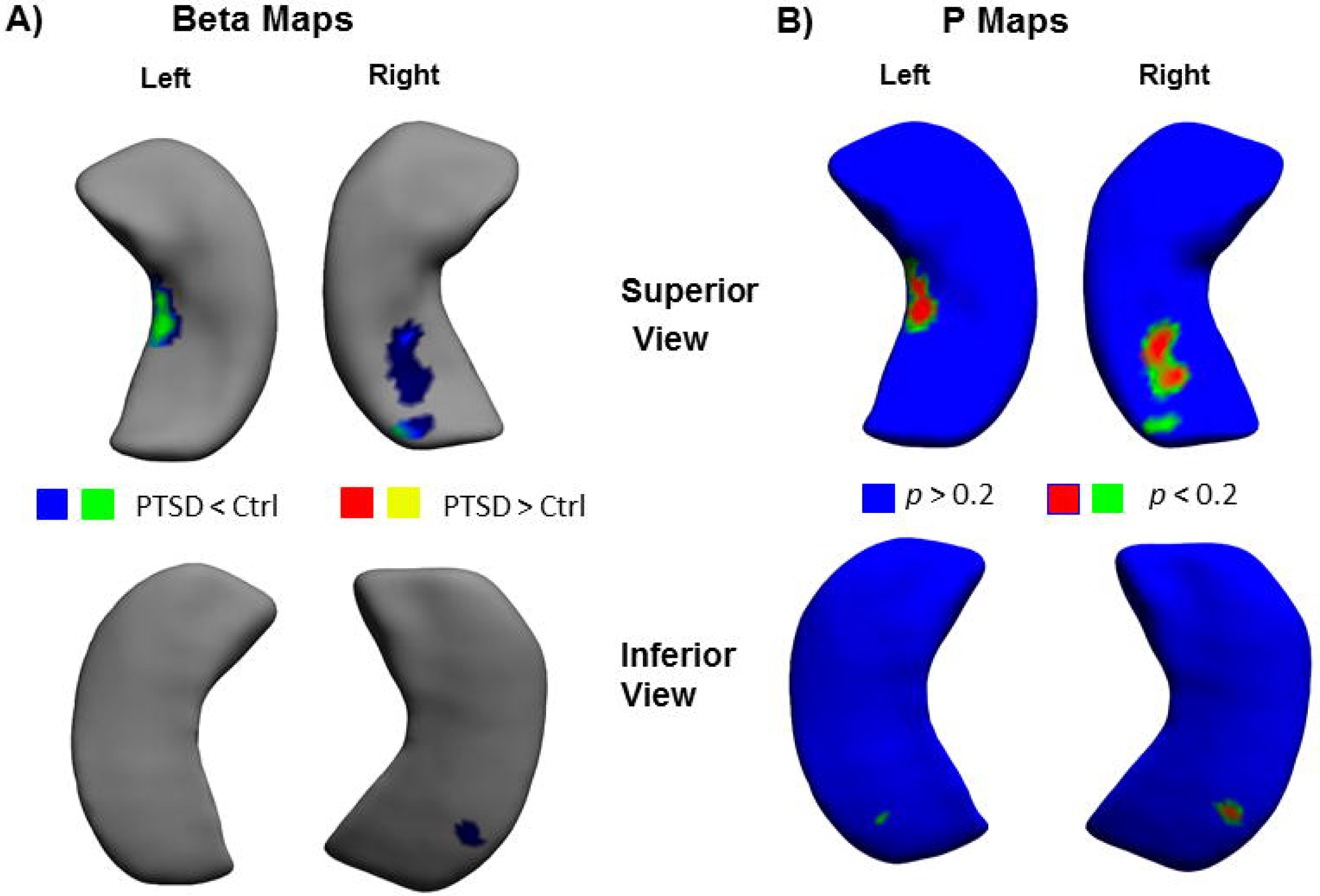
Shape analysis. The results of 3D shape analysis at 2,502 vertices show with FDR correction at *p* < 0.2 significance level. (A) β-map show PTSD < Control and PTSD > Control **(B)** *p*-map.

### Effect of Scanner

There was no systematic difference in the number of cases and controls across scanner [X^2^=4.48; p = .11]. Subfield findings between scanners did not differ significantly based on PTSD diagnosis when controlling for age. The interaction of PTSD and scanner type was non-significant for L-CA1 [F_278,2_=1.29, p=0.28], L-CA3 [F_278,2_=1.40, p=0.25], and R_CA3 [F_278,2_=1.58, p=0.21].

## DISCUSSION

In this study we examine PTSD-associated differences in the volume of hippocampal subfields and the shape of the hippocampus. Our study demonstrates smaller volume in the L-CA1, L-CA3, and R-CA3 in patients with PTSD. Among the present findings, only L-CA1 had a non-trivial effect size, whereas L-CA3 and R-CA3 had very small effect sizes. While trauma-exposure was significantly higher in the PTSD group than the trauma-exposed control group, we found no evidence to support that between-group differences in subfield volumes were correlated to severity of trauma exposure. We found an inverse correlation of the L-DG subfield with age in the combined group, and with L-CA1, and L-DG in the PTSD group, but this relationship was unaffected by PTSD diagnosis. No PTSD-associated differences in hippocampal shape were detected with vertex-based morphometry. Compared to previously published reports of hippocampal subfield volume in PTSD [7], our study used a sample size that is over 2.5 times larger and employed more advanced methods for segmentation of 12 hippocampal subfields based on an atlas constructed from ultra-high resolution MRI scans of postmortem brains.

Lesion studies in model animal systems that introduced neurotoxic lesions in either CA1 or CA3 prior to extinction training eliminated the context dependence of extinguished fear. However lesions of CA1 or CA3 placed after extinction training found that only CA1 lesions impaired context dependence of extinction [19]. Thus, hippocampal CA1 neurons may play an important role in conditioned fear and extinction [20]. Insofar as contextual fear extinction is a realistic model of PTSD and the assumption that reduced CA1 volume implies a corresponding function in humans, our finding is consistent with the clinical features of PTSD, specifically the presence of prominent re-experiencing symptoms that persist in contexts without threat.

Hippocampal subfields CA1–3 are altered by experimental stress in animal studies [34] and consistent with human behavioral findings of the preferential impact of early life maltreatment stress in at-risk families [35]. One consequence of this type of stress is corticosteroid-mediated impairment of learning and memory including the formation of false memories [36]. The basolateral amygdala, which is critical to associative emotional learning, projects directly and indirectly through the entorhinal cortex to CA1 [37]. Pavlovian fear conditioning increases synchronization of the basolateral amygdala with CA1 and facilitates consolidation of declarative memory. Whereas repeated exposure to acute restraint stress produced hypertrophy of spine density in the basolateral amygdala and increase in anxiety-like behavior in the mice, it resulted in long term depression and atrophy in CA1 [38; 39]. Preclinical research in rats has shown that the CA1 subfield is involved in context-specific memory retrieval after extinction [19]. Ventral CA1 is enriched is enriched in anxiety cells that are activated by anxiogenic environments and required for avoidance behavior in mice [40]. Amygdala-hippocampal interactions are important for both non-declarative and declarative forms of emotional memory. Thus, our finding of lower CA1 volume in PTSD is consistent with the well-designed carefully controlled experiments in animals. In PTSD, the basolateral amygdala and the hippocampus function to control effects of emotion and arousal on consolidation of memories that include spatial or contextual cues about the environment [41].

To interpret the left lateralization of the present CA1 findings in PTSD, we turn to evidence from humans to adequately consider the unique role of language and verbal memory, which are lateralized to the left hippocampus [22]. Kerchner and colleagues [42] found that neuronal volume in CA1 alone was associated with episodic memory for verbal, visuospatial, and logical information. This association of CA1 with episodic memory was left lateralized and not found in relation to CA3 or DG. Disruption in verbal memory is a prominent feature of PTSD that has been demonstrated with neurocognitive testing and the inability to recall granular details of traumatic experiences is consistently observed in clinical encounters [36; 43; 44]. These memory disruptions are correlated with left lateralized decrements in hippocampal activation [36; 45]. Thus, converging evidence from behavioral, neuropsychological, functional imaging, volumetry, and lesion studies support a compelling rationale for left-lateralized CA1 alterations. Teicher and collegues reported lower volume of CA1, CA2/CA3, fimbria, presubiculum, subiculum, and CA4/DG in adults who experienced maltreatment as children, with 7% of the sample diagnosed with PTSD. However, there are a number of methodologic differences between the present study and Teicher et al. FreeSurfer 5.3 is based on an anatomically incorrect atlas [9], notably for CA1. In studies of PTSD that employed manual subfield segmentation, Wang et al reported lower volume in CA3, whereas Mueller et al found no subfield differences associated with PTSD. Several factors for divergent findings in previous studies may include lack of power, differences in military service, education, age, race, and socioeconomic status between samples. Small effect sizes calculated for the present results show that previous studies conducted with a sample sizes of n=36 [6] and n=85 [5] were underpowered. While our findings recapitulate the age associated volume decrease in CA1, we did not find a significant interaction aging and PTSD for CA3 volume as reported by Wang et al., [6].

As expected, directly comparing results across studies poses several challenges because of methodologic heterogeneity. The present segmentation with FreeSurfer v6.0 was fully automated, with the exception of quality checks performed on 12 segmented labels. However, Wang et al. and Mueller et al. employed manual segmentation, which may be able to overcome image artifacts sometimes overlooked by automated techniques. However, recent developments in automated methods take advantage of generative models for multi-atlas image segmentation that do not rely on the intensity of the training images. The, generative approaches used in FreeSurfer v6.0 and SPM have bias field estimation integrated into the model and are therefore very robust to bias fields. However we actually applied the FreeSurfer segmentation with the bias field correction turned off because the *norm.mgz* is already bias field corrected and moreover the bias field around the hippocampus is negligible. Noise is also integrated into the framework by modeling Gaussian distributions. An explicit model for motion is challenging to implement because it would need to contend with a variety of pulse sequences. Thus, severe motion can indeed derail the algorithm, but in practice it works very well [46]. Typically, the spatially varying tissue priors used by automated segmentations are produced by humans and training sets used by machines are also initially classified by humans [47].

FreeSurfer v6.0 combines *ex-vivo* and *in-vivo* scans with the former acquired on 15 postmortem brains scanned at 7-Tesla with on average 130-µm isotropic resolution, whereas v5.3 used in vivo atlas from five cases acquired at 9-fold lower resolution of 380-µm3 resolution and manual segmentation studies were acquired at 4T with 180-fold lower resolution of 400 ×; 500 µm in-plane and 2000-µm through-plane resolution. Although the landmarks chosen to delineate the subfield boundaries on the high resolution 7T *ex vivo* images were defined based on knowledge derived from histological exams, the accuracy of the resulting *ex vivo* labels was not confirmed by a histological exam in these specimens.

We conducted analyses with and without the inclusion of ipsilateral whole hippocampal volume as a covariate. Our reasoning for including ipsilateral whole hippocampal volume covariate is based on the fact that CA1 volume is highly correlated with whole hippocampal volume, i.e. individuals with smaller hippocampi will have smaller CA1 in much the same way that individuals with a small brain (TIV) will have a small hippocampus. This correlation is strong between L-CA1 and L-whole hippocampal volume in the control group (R^2^=0.8647) and in the PTSD group (R^2^=.8679). Thus, including the covariate for whole hippocampal volume is more effective at accounting for sources of variance, in particular for individual differences in size for both control (R^2^=0.3565) and PTSD (R^2^=0.3197) groups. While it is possible that smaller overall hippocampal volume in PTSD might partially mask the association between PTSD and subfield volume, in fact, it appears that this source of variance contributes more to the model than diagnostic status. Additional arguments for including ipsilateral whole hippocampal volume over TIV include (1) TIV does not account for laterality effects since it encompasses both hemispheres unlike ipsilateral whole hippocampal volume which is hemisphere-specific, and (2) the subfield volume is highly correlated with ipsilateral whole hippocampal volume because the FS v6.0 subfield segmentation relies heavily on atlas priors as compared to specific anatomical landmarks. By contrast, not including ipsilateral whole hippocampal volume answers a different but interesting question, which is whether subfield volume is associated with PTSD but without regard to the volume of the whole hippocampus. The association of the ‘absolute subfield volume’ to PTSD is also important because it does not presuppose the role of whole hippocampal volume or the association between PTSD and whole hippocampal volume. A thorough and thoughtful treatment of multiple regression modeling with multiple collinear variables is provided by Friedman and Wall [48] and also by Wurm and Fisicaro [49].

An estimation of statistical power for the largest published study (n=97) [7] to detect group differences in CA1 volume based on the effect size we observed (0.21), reveals only 55.3% power to reject the null hypothesis or a 44.7% chance of a false positive. By comparison, the present sample size (n=282) based on the same effects size (0.21) attains 94.9% power to reject the null or a 5.1% chance of a false positive. Clearly the results of previously published studies are concerning for reproducibility with a false positive chance approaching 50%.

## Limitations

An important limitation in interpreting our finding of diminished CA1 volume is the lack of correlative behavioral measures memory. Lacking such data, only indirect and tenuous associations can be made between PTSD diagnosis and memory disturbance, which can be inferred from symptoms of recurrent intrusive trauma memories and trauma reliving, or by invoking prior reports that demonstrate PTSD is associated with disrupted fear memory [50]. Nonetheless, we feel that the outcome of the present study could generate future work to test these predictions.

FreeSurfer v6.0 segmentation was qualitatively compared with manual segmentation on a publicly available dataset of high resolution T1/T2 images [27] and quantitatively compared with the ADNI dataset, which involved discriminating MCI and normally aging elderly subjects. FreeSurfer compares favorably to manual segmentation based on extensive testing of FreeSurfer v6.0 as described by Iglesias et al [10].

A notable limitation of the FreeSurfer 6.0 scheme for parcellation is that it depends heavily on a shape prior from an atlas composed of 15 ex vivo ultra-high resolution images acquired at 7T and hence delineates structures that cannot be discerned visually on in vivo 1-mm isotropic images with sufficient clarity to accurately parcellate 12 subfields. This difference of approach is a concern because the algorithm sometimes relies on the shape priors for labeling rather than contrast information detected in the image being segmented [10]. The lack of sufficient contrast features from the in vivo data makes robust quality control a challenge even by the most expert manual rater.

Our results lacked significant shape findings related to PTSD after applying the correction for multiple comparisons. While the corrections are applied to control Type 1 error, the procedure invariably comes at the cost of inflating Type II error. On the other hand, we imposed no correction on the volumetry results for CA1 because of a priori evidence implicating CA1 in PTSD, which eliminated any risk of Type II error. Thus, the first possible explanation relates to the application of multiple comparison correction in the shape analysis but no correction in the volumetry analysis of CA1. It does appear that the uncorrected shape results show differences in the area of CA1. The second explanation is that shape differences are just below the threshold that is deemed significant (e.g. p > 0.05). However, it is possible that the subthreshold vertex-based differences accumulate to produce a volume difference that is significantly different between groups. This is all the more likely given that the effect size of volume differences was small even for the CA1, which was the largest effect size among the subfields. The third possibility is simply some combination of the first two.

A limitation is our use of three different scanners (two GE 3T and one Philips 3T). We have previously analyzed FreeSurfer v6.0 output from different scanners with evidence that scanner heterogeneity provides robust and consistent results for hippocampal segmentation [4] and hippocampal subfield segmentation [25].

## Conclusions

Our results provide robust evidence of an association between smaller hippocampal L-CA1 volume and PTSD. Lower CA1 volume in PTSD is consistent with established research in mice that demonstrates atrophy in CA1 and increased anxiety-like behavior results from repeated exposure to acute stress. Further work on hippocampal subfield plasticity in PTSD with longitudinal studies will be important to elucidate the role of subfield alterations on learning and memory impairments in PTSD, particularly associative fear learning, extinction training, and the formation of false memories.

## ACKNOWLEDGMENTS

This research was supported by the U.S. Department of Veterans Affairs (VA) Mid-Atlantic Mental Illness Research, Education, and Clinical Center (MIRECC) core funds. Dr. Morey also received financial support from the VA Office of Research and Development (5I01CX000748-01, 5I01CX000120-02). Additional financial support was provided by the National Institute for Neurological Disorders and Stroke (R01NS086885-01A1). Juan Eugenio Iglesias is supported by a Starting Grant from the European Research Council (project no. 677697: “BUNGEE-TOOLS”). Emily Dennis is supported by a grant from the NINDS (K99 NS096116). Emily Dennis, Christopher Whelan, Boris Gutman, Neda Jahanshad, and Paul Thompson are also supported by NIH grants to Paul Thompson: U54 EB020403 (BD2K), R01 EB008432, R01 AG040060, and R01 NS080655. Finally, we could not have conducted this study without the willing participation of Veterans.

---

**Supplemental Figure 1.**
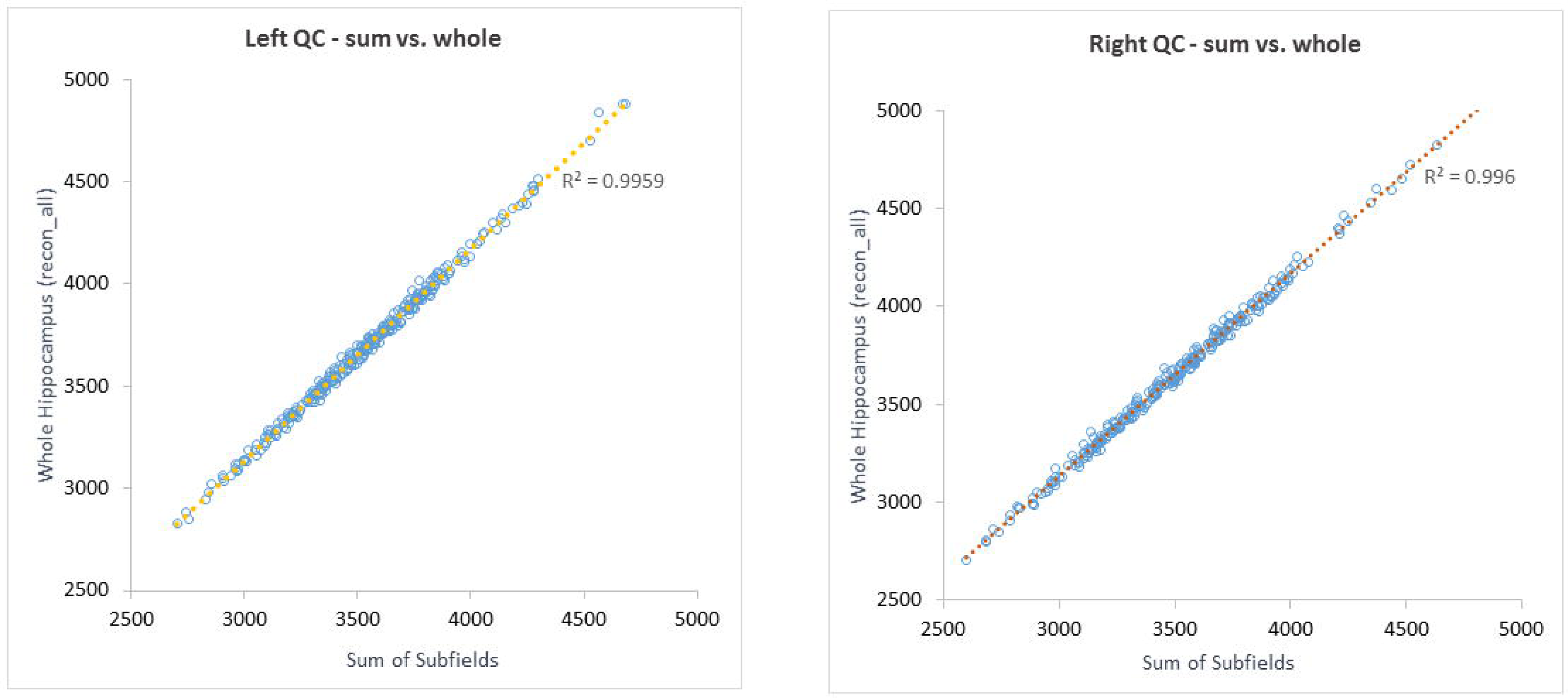

**Supplemental Figure 2.**
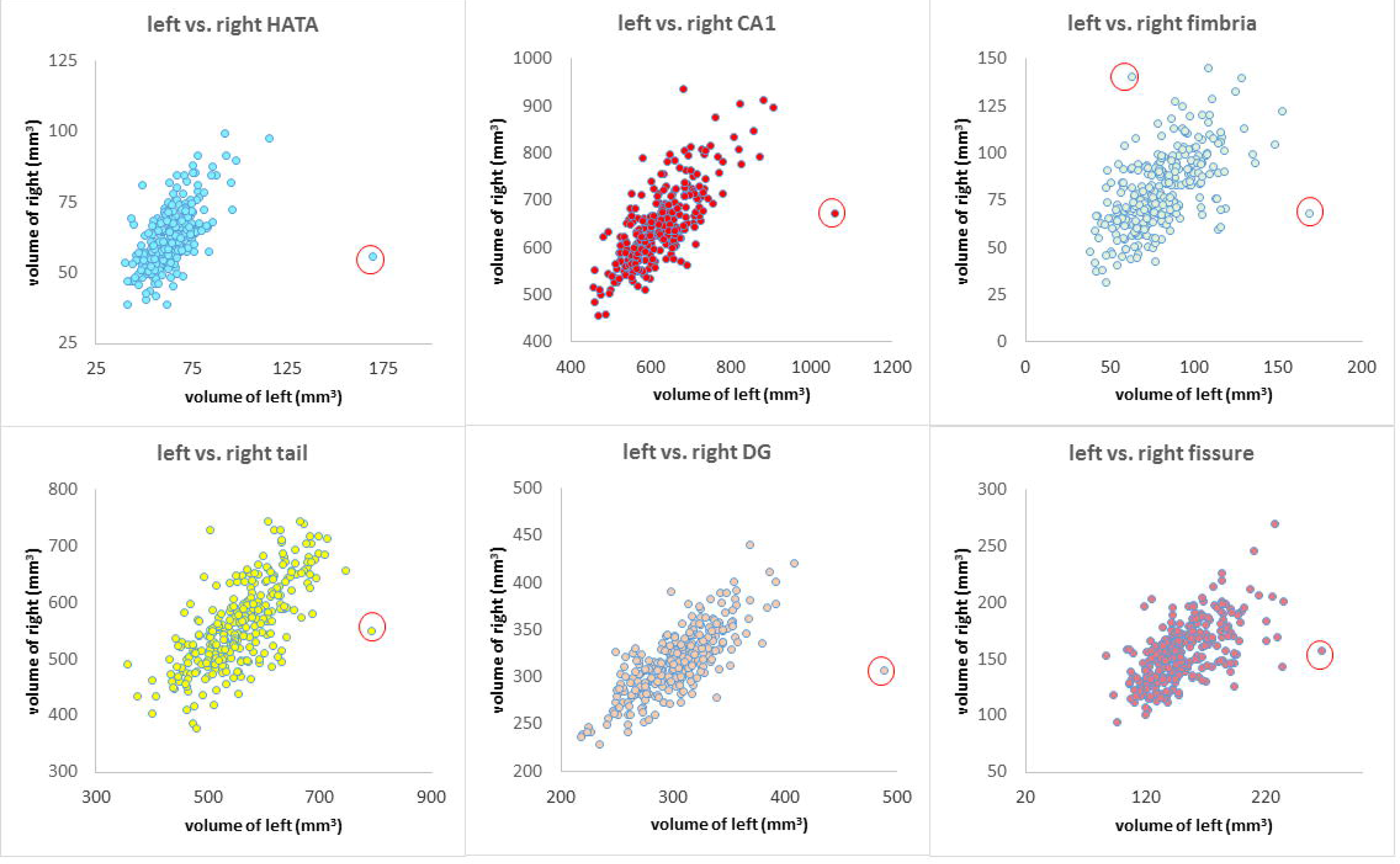
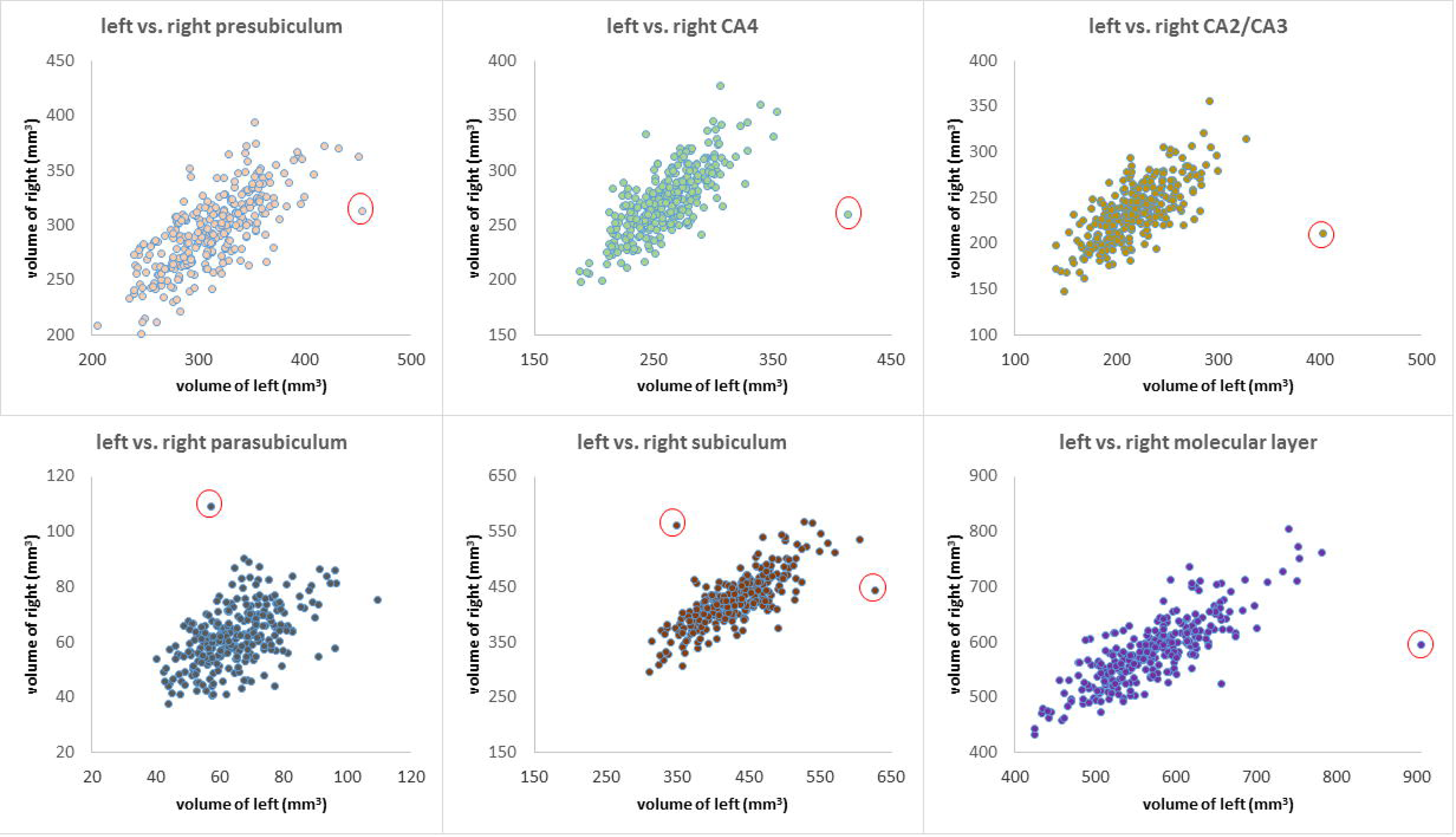

**Supplemental Figure S3.**
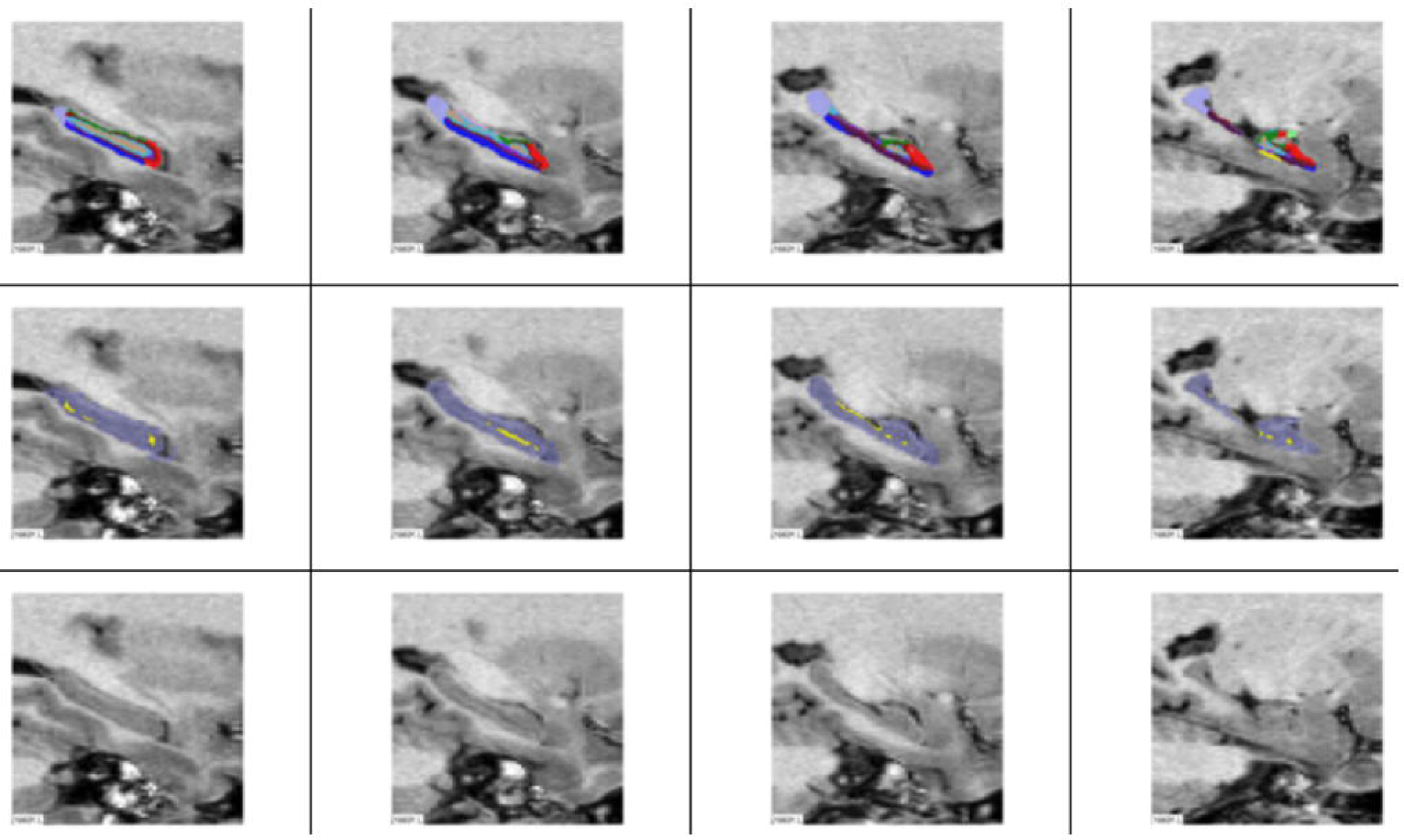

